# Reward boosts neural coding of task rules to optimise cognitive flexibility

**DOI:** 10.1101/578468

**Authors:** S. Hall-McMaster, P.S. Muhle-Karbe, N.E. Myers, M.G. Stokes

## Abstract

Cognitive flexibility is critical for intelligent behaviour. However, its execution is effortful and often suboptimal. Recent work indicates that flexible behaviour can be improved by the prospect of reward, which suggests that rewards optimise flexible control processes. Here we investigated how different reward prospects influence neural encoding of task rule information to optimise cognitive flexibility. We applied representational similarity analysis (RSA) to human electroencephalograms, recorded while female and male participants performed a rule-guided decision-making task. During the task, the prospect of reward varied from trial to trial. Participants made faster, more accurate judgements on high reward trials. Critically, high reward boosted neural coding of the active task rule and the extent of this increase was associated with improvements in task performance. Additionally, the effect of high reward on task rule coding was most pronounced on switch trials, where rules were updated relative to the previous trial. These results suggest that reward prospect can promote cognitive performance by strengthening neural coding of task rule information, helping to improve cognitive flexibility during complex behaviour.

**Significance Statement:** The importance of motivation is evident in the ubiquity with which reward prospect guides adaptive behaviour and the striking number of neurological conditions associated with motivational impairments. In this study, we investigated how dynamic changes in motivation, as manipulated through reward, shape neural coding for task rules during a flexible decision-making task. The results of this work suggest that motivation to obtain reward modulates encoding of task rules needed for flexible behaviour. The extent to which reward increased task rule coding also tracked improvements in behavioural performance under high reward conditions. These findings help inform how motivation shapes neural processing in the healthy human brain.

## Introduction

Flexible cognitive control is critical to human intelligence. When vying to win a card game, we can use arbitrary rules to play the best hand. When navigating a new city, we can apply navigation rules to sensory input from the world around us to arrive at the next tourist attraction. Controlled processes requiring flexibility result in slower performance, compared to behaviours that require less flexible processing (Kleinsorge & Rinkenauer, 2012; Shen & Chun, 2011). However, this can be improved by motivational factors, such as the prospect of reward for speed and accuracy (Braem & Egner, 2018). When the prospect of reward is high, performance improves on flexible rule-based tasks (Etzel et al., 2015; Kleinsorge & Rinkenauer, 2012), suggesting that reward might optimise cognitive flexibility by fine-tuning rule-based neural coding patterns.

The existing neuroimaging literature provides support for this perspective, indicating that reward prospect leads to stronger recruitment of frontoparietal brain regions implicated in cognitive control (Parro, Dixon, & Christoff, 2018). When reward cues are presented at the beginning of each trial, activity in these regions is typically enhanced prior to the onset of a target stimulus, suggesting that proactive control mechanisms contribute to reward-induced performance benefits (Engelmann et al., 2009; Krebs et al., 2011; Padmala & Pessoa, 2011). Critically, a recent study combining fMRI with pattern classification methods found that frontoparietal activity prior to target onset encoded abstract task rules with greater fidelity on reward trials than on no-reward trials (Etzel et al., 2015). The extent to which reward prospect enhanced the decodability of task rules also mediated behavioural benefits, suggesting that performance improvements may result from reward-driven tuning in cognitive control processes that prioritise task-relevant processing.

This proposal has considerable theoretical appeal. Yet, it is unknown whether reward prospect differentially influences rule coding in situations that require more or less flexible processing. The present study therefore aimed to characterise how changes in reward influence neural coding of task rule information when rules must be updated. To do so, we developed a behavioural task that involved switching between task rules, as reward prospect was manipulated. We then applied representational similarity analysis (RSA; Kriegeskorte, Mur, & Bandettini 2008) to human electroencephalograms to examine changes in neural coding as a function of reward prospect. Based on Etzel et al. (2015), we predicted that task rule coding would be greater with high reward prospect and that the extent of this increase would track improvements in behavioural performance. However, the critical contribution of this work was to consider how reward prospect would impact rule coding when flexible processing was needed. Based on prior work showing selective reward enhancements on rule switching (Shen & Chun, 2011; Kleinsorge & Rinkenauer, 2012), we reasoned that task rule coding should be greatest on high reward switch trials, where task rules must be updated. Switch trials involve the most interference between rule codes and thus increased neural separation between them should support flexible rule updating (Waskom, Kumaran, Gordon, Rissman, & Wagner, 2014). Reward prospect could therefore optimise flexible processing by helping to increase neural dissimilarity between rules, when they come into conflict. RSA is well suited to test this prediction because it provides a sensitive index of reward-driven changes in neural dissimilarity, while allowing clean separation of overlapping task variables in the lead up to adaptive behavioural responses.

To summarise the main results, we found that high reward prospect produced significant performance improvements in accuracy and reaction time (RT). Consistent with the view that reward prospect increases proactive cognitive control, we found a significant increase in neural coding for task rules under high reward conditions prior to the onset of a target stimulus. The average difference in rule coding between reward conditions during this period also correlated with RT improvements. Especially striking, reward’s effect on rule coding was most pronounced in situations requiring the most flexible processing, when task rules were updated relative to the previous trial.

## Materials and Methods

### Participants

We set a target sample size of 30 participants. During recruitment three participants were excluded, one due to a corrupt EEG recording and two due to excessive artefacts that led to the rejection of more than 120/650 trials. We therefore collected three more participants to reach the 30 participant target. The final sample were between 18 and 35 years of age (mean age = 23, 19 female), with normal or corrected-to-normal vision, who reported no history of neurological or psychiatric illness. Participants received £8 per hour or course credit for taking part and could earn up to £10 extra for their performance. This study was approved by the Central University Research Ethics Committee at the University of Oxford and all participants signed informed consent before taking part.

### Materials

Stimuli were presented on a 22-inch screen with a spatial resolution of 1280 × 1024 and refresh rate of 60Hz. Stimulus presentation was controlled using Psychophysics Toolbox-3 (Kleiner, Brainard & Pelli, 2007) in MATLAB (MathWorks, version R2015b). Reward cues and feedback shown during the task were presented in size 30 Arial font. Task cues and target stimuli had approximate visual angles of 2.52° (100×100 pixels) and 1.26° (50×50 pixels) respectively, with visual angles calculated based on an approximate viewing distance of 60cm. F and J keys on a standard QWERTY keyboard were used to record left and right hand responses. EEG data were recorded with 61 Ag/AgCl sintered electrodes (EasyCap, Herrsching, Germany), a NeuroScan SynAmps RT amplifier, and Curry 7 acquisition software (Compumedics NeuroScan, Charlotte, NC). EEG data were pre-processed in EEGLAB (Delorme & Makeig, 2004, version 14.1.1b), behavioural analyses were performed using JASP (https://jasp-stats.org, version 0.8.1.3) and EEG analyses were performed in MATLAB (MathWorks, version R2015b) using the EEGlab and Fieldtrip toolboxes as well as custom scripts.

### Code Accessibility

Task and analysis code, as well as raw and pre-processed data will made openly available, following acceptance of the manuscript.

### Experimental Design and Statistical Analysis

In this task (Figure 1), participants’ overarching goal was to gain as many points as possible. To do so, participants categorized bi-dimensional target stimuli based on their colour (yellow vs blue) or shape (square vs circle). On each trial, only one feature dimension of the target (colour or shape) was relevant to gaining points, while the other feature served as an irrelevant distractor. The relevant feature dimension was signalled through a visual task cue prior to target onset. In addition to a single relevant feature dimension, each trial offered a high or low reward magnitude for making a correct response. This was signalled to participants at the beginning of each trial by a single pound sign (low reward: 5-10 points) or three pound signs (high reward: 50-100 points).

**Figure 1.**
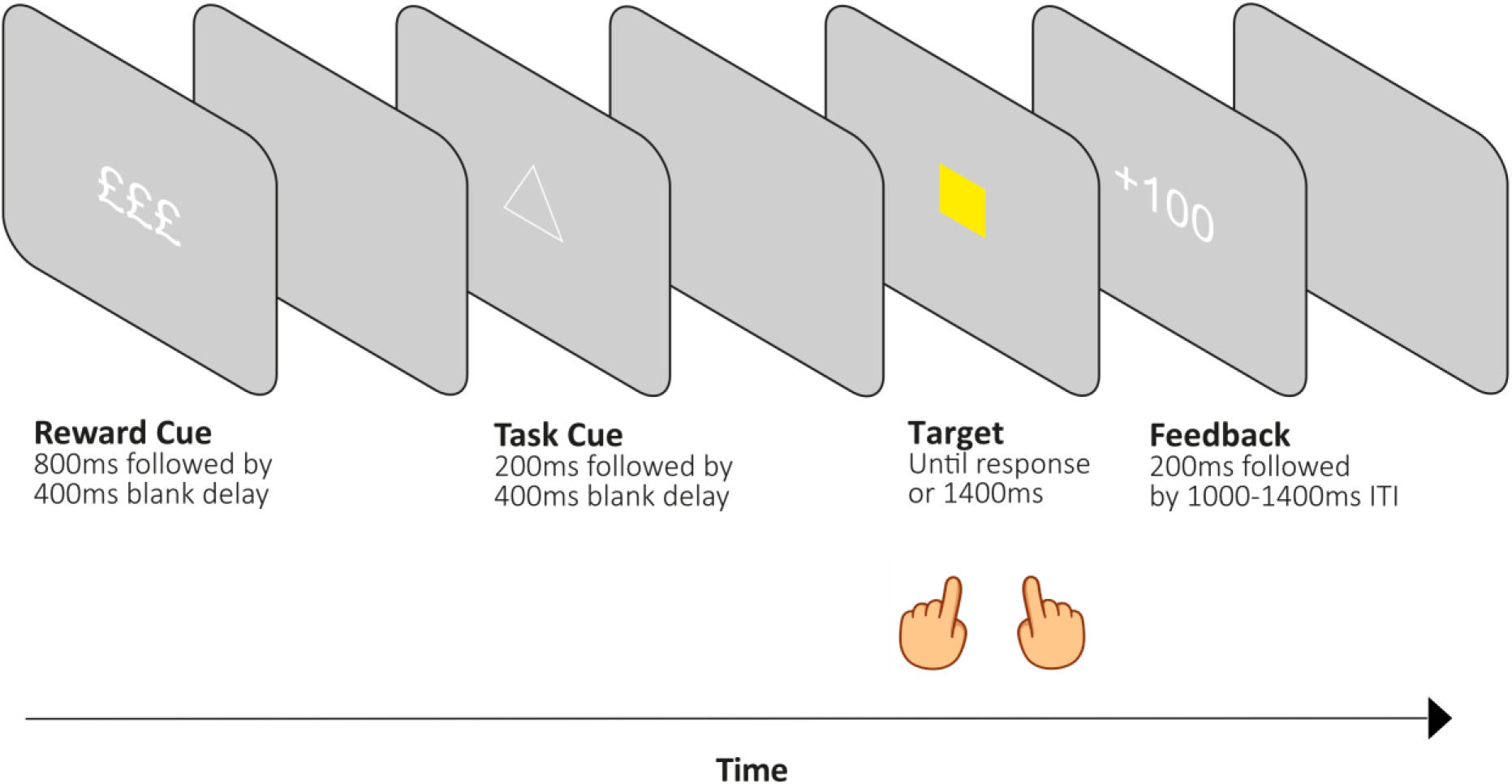
Task design. On each trial, a high or low reward cue was presented followed by a blank delay. A task rule cue was then presented, indicating whether participants should respond to the upcoming target based its colour or shape. Following a second blank delay, a bi-dimensional target (a coloured shape) appeared until a response was given or for a maximum duration of 1400ms. This was followed by feedback (based on accuracy and reaction time) and a variable inter-trial-interval.

The experimental sequence consisted of a reward cue, task cue, target, feedback screen and an inter-trial interval (ITI). The reward cue (£ or £££) was first presented for 800ms, followed by a 400ms delay. The task cue (one of four possible abstract shapes) was then presented for 200ms. The mapping of cues to tasks was counterbalanced between participants. Task cue offset was followed by a 400ms delay. The target (a yellow square, blue square, yellow circle or blue circle) was then presented and remained on screen until a response was given or for a maximum of 1400ms. If the active task rule was colour, the correct response mapping was ‘f’ or ‘j’ for yellow or blue targets respectively. If the active rule was shape, the correct response mapping was ‘f’ or ‘j’ for square or circle targets respectively. The response phase was followed by feedback lasting 200ms. An incorrect response or omission resulted in feedback showing ‘+0’ points. A correct response resulted in feedback showing ‘+X’, where X was a value within the high or low reward point ranges, the precise value of which was determined by RT. More specifically, RT criteria for different points were initialised so that responses faster than 400ms, 600ms, 800ms, 1000ms, 1200ms and 1400ms earned 100/10, 90/9, 80/8, 70/7, 60/6 and 50/5 points on high and low reward trials respectively. For correct trials, the current trial RT was added to an array for its reward condition. When each array contained more than six values, individualised points criteria were calculated for that condition and were calculated again every time a new value entered the array. The individualised points criteria followed criteria outlined by Shen & Chun (2011), in which the most (to least) points are rewarded for correct responses faster than 95%, 80%, %65, %50 and 35% of the median RT. The trial concluded with a randomly selected ITI duration, drawn from a uniform distribution with values of 1000, 1100, 1200, 1300 or 1400ms. Participants were trained to reach a criterion of 70% accuracy before completing 10 experimental blocks of 65 trials. Excluding the first trial in each block, equal numbers of reward cues, task cues, stimuli and ITI durations were presented within each block. Presentation was pseudo-randomised to ensure trials were balanced based on task, target-congruency and task sequence for each reward condition. Target congruency refers to whether task-relevant and irrelevant target features are mapped to the same (congruent) or different (incongruent) response hands. Task sequence refers to whether the task rule was different from the previous trial (switch trial) or the same as the previous trial (repeat trial).

#### Behavioural measures

Our main dependent measures were reaction time (RT) and the proportion of correct responses (accuracy). We calculated median RTs for the respective design cells to account for the skewness of RT distributions (see Ratcliff, 1993). For all RT analyses, we only included trials in which the current and previous trials were correct to mitigate potential effects of post-error slowing, in which participants tend respond more slowly after making an error (Dutilh et al., 2012; Notebaert et al., 2009). For accuracy analyses, we included all trials in which a response was made within the 1.4s response window. Behavioural data were analysed using 2×2 repeated measures ANOVAs with factors of reward and task sequence. The task sequence factor consisted of switch trials, where the task rule was different from the previous trial, and repeat trials, where the task rule was the same as the previous trial. In addition, we used a paired sample t-test for normally distributed data and a Wilcoxon signed-rank test for data showing significant deviation from normality, to compare performance on task-switch trials and cue-switch trials. This analysis served as a behavioural control. It has been argued that switch costs could arise simply from processes related to processing the cue stimulus, rather than changes in task rule per se (Logan & Bundesen, 2003; Logan & Schneider, 2006). Comparing trials where both task cue and task rule changed, with trials where the cue changed but the rule stayed the same allowed us to delineate the extent to which switch costs were driven by changes in task rules and task cues. More specifically, if the effect of rule switching on behavioural performance was primarily driven by rule changes, we should see greater performance reductions on trials where the rule is changed relative to trials where the rule is repeated but the cue is changed.

#### EEG pre-processing

EEG data were down-sampled from 1000 to 250Hz and filtered using 40Hz low-pass and 0.01Hz high-pass filters. For each participant, channels with excessive noise were identified by visual inspection and replaced via interpolation, using a weighted average of the surrounding electrodes. Data were then re-referenced by subtracting the mean activation across all electrodes from each individual electrode at each time point. Data were divided into epochs from −1 to +5 seconds from the onset of the reward cues. Epochs containing artefacts (such as muscle activity) were rejected based on visual inspection. In the final stage of pre-processing, data were subjected to an Independent Component Analysis (ICA). Structured noise components, such as eye blinks, were removed, resulting in the data set used for subsequent analyses. Prior to each analysis, data were z-scored over the trial dimension and baseline-corrected using a time window of 200ms to 50ms prior to the trial event of interest (e.g. cue or target presentation).

#### EEG analyses

We used RSA (Kriegeskorte et al., 2008) to investigate how reward prospect influenced neural coding for different kinds of task-relevant information. There were two main advantages of using RSA to address this question. First, multivariate approaches leverage pattern information that would normally be averaged out in univariate analyses. This makes multivariate methods more sensitive to effects based on distributed patterns of neural activity (Kriegeskorte, Goebel, & Bendettini, 2006). Second, RSA allowed us to examine multiple, overlapping neural codes. In particular, the combination of RSA with models for different task variables provided a powerful method for separating neural coding of task rules, relevant and irrelevant target features, as well as motor responses. This ability to separate overlapping neural activity would be difficult to achieve with more traditional EEG methods, such as the analysis of event-related potentials.

In addition, we selected EEG for its high temporal resolution. Constraints on the temporal resolution of fMRI can make it challenging to isolate task rule coding from subsequent perceptual processing because the slow hemodynamic response can make it difficult to pinpoint effects in time and distinguish sustained anticipatory activity from transient stimulus-evoked responses. In contrast, high temporal resolution methods such as electroencephalography are needed because their ability to distinguish rapid stimulus-evoked dynamics makes them ideal for isolating the effects of reward on task rule coding from subsequent neural coding patterns.

The logic of our approach was to characterise neural coding patterns elicited by different trial conditions and test whether reward prospect led to more distinct task representations for the two tasks being performed (colour vs shape judgements; Etzel et al., 2015; Westbrook & Braver, 2016). We were especially interested in the effect of reward on proactive control mechanisms, wherein goal-relevant information is encoded in preparation for upcoming task demands. This stands in contrast to reactive control processes, which serve to resolve task demands after their detection (Braver, 2012). Evidence of reward-modulated proactive control in our design would be seen as a difference in rule coding between high and low reward conditions, prior to target onset. Moreover, the difference in rule coding during this period should also be associated with improvements in behavioural performance (Etzel et al., 2015). Building on the results of Etzel et al. (2015) a central goal in the present work was to examine the effect reward prospect on rule coding during switch trials, when rules required flexible updating. Here we reasoned that switch trials involve the greatest interference between rule representations and thus reward prospect could improve flexible cognitive control (Kleinsorge & Rinkenauer, 2012; Shen & Chun, 2011) by increasing neural dissimilarity between rule codes. As a secondary aim, we performed theory-driven analyses that tested whether high reward lead to stronger neural coding for task-relevant compared to irrelevant target features, and whether relevant feature prioritisation was associated with improved performance (Pessoa, 2017). Finally, we performed exploratory analyses that tested whether high reward modulated motor response coding and whether reward-induced changes in task coding were associated with downstream changes in sensorimotor processing. All analyses used data from all 61 EEG channels. While this limits inference about regional sources of neural activity, it has the advantage of including all available data without any assumptions about source localization. Incorrect and omission trials were excluded from EEG analyses.

##### Neural coding across the trial

For each participant, trials were divided into conditions based on reward condition (low, high), as well as task-relevant and irrelevant target features. Dividing the trials this way also implicitly divided trials by task. If the task-relevant target feature was yellow, for example, then the task must have been to judge the target colour. We then averaged trials in each condition to get an array with channels x time points x conditions. To measure neural dissimilarity between conditions, we used Mahalanobis distances (MDs). This distance metric shows similar reliability to correlation distance measures (Walther et al., 2016) but explicitly takes covariance into account, making it well suited to EEG data where channel values tend to be highly correlated. For EEG data, the MD between two conditions is computed at each time point, using two key pieces of information. The first is the difference in topographies between two conditions. A topography is a vector of 61 channel values that has been averaged across all trials in a condition, for a specific time point (Figure 2A). The second key piece of information is the channel covariance matrix. This is computed using a matrix of trials x channels. For this study, we used within-condition error to compute the channel covariance matrix (Walther et al., 2016). This meant that prior to computing the covariance matrix, trials in the trial x channel matrix were mean-centered by subtracting the mean topography for each condition from all trials within that condition. The covariance calculation also used shrinkage estimator (Ledoit & Wolf, 2004), which has the effect of down-weighting noisy covariance estimates. Using this information, the MD is formally computed as: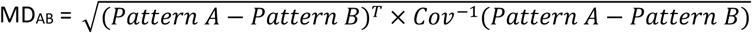, where Pattern A-Pattern B is the difference between topographies, T is the transpose and Cov-1 is the inverse of the channel covariance matrix. This calculation is also presented in Figure 2. MDs were calculated between all topographies for the sixteen conditions, separately at each time point. This procedure yielded a 16 × 16 representational dissimilarity matrix (RDM) of multivariate condition distances for each time point and participant. We then constructed a set of five 16 × 16 model RDMs to capture neural coding patterns related to different task variables. These variables included reward coding, task coding, task-relevant feature coding, task-irrelevant feature coding, and motor coding. The logic of all models was to place zeroes in cells of a 16×16 matrix where conditions matched on the variable of interest and ones in all remaining cells. For example, the task coding model was a 16×16 matrix containing zeros in cells where two conditions involved the same task (e.g. both colour judgements) and ones in cells where two conditions involved different tasks (e.g. shape vs colour judgements). The task-relevant feature model was a 16×16 matrix containing zeros in cells where two conditions had the same task-relevant target feature (e.g. both yellow on colour trials) and ones in cells where two conditions had different task-relevant features (e.g. yellow vs blue or yellow vs square). Data RDMs (not z-scored and averaged for illustration) and model RDMs for all task variables are presented in Figure 3. For analysis, data and model RDMs were z-scored. As RDMs are symmetric, the upper triangular portion of each matrix was transformed into a vector. The resulting data and model distance vectors were then entered into a multiple regression analysis that was conducted at each time point (4ms apart after down-sampling). The data-derived distance vector was the dependent variable and the model-derived distance vectors were independent variables. The regression also included a constant to model the intercept of the linear regression equation. This led to the following general linear model (where DV stands for distance vector): *Data DV* = *β*0(*intercept*) + (*β*1 *x reward DV*) + (*β*2 *x task DV*) + (*β*3 *x task relevant feature DV*) + (*β*4 *x task irrelevant feature DV*) + (*β*5 *x motor response DV*) + *ε*(*residual error*). The regression procedure was performed three times, once with a baseline window prior to reward cue onset, once prior to task cue onset and once prior to the target onset. Running the regression procedure at each time point produced a time course of regression coefficient estimates, one for each coding model. We refer to coefficient estimates as ‘model fit to neural dissimilarity matrix’ in subsequent figures. We interpret the magnitude of these coefficient estimates to reflect the magnitude of neural dissimilarity/neural coding for each of the task variables across time.

**Figure 2.**
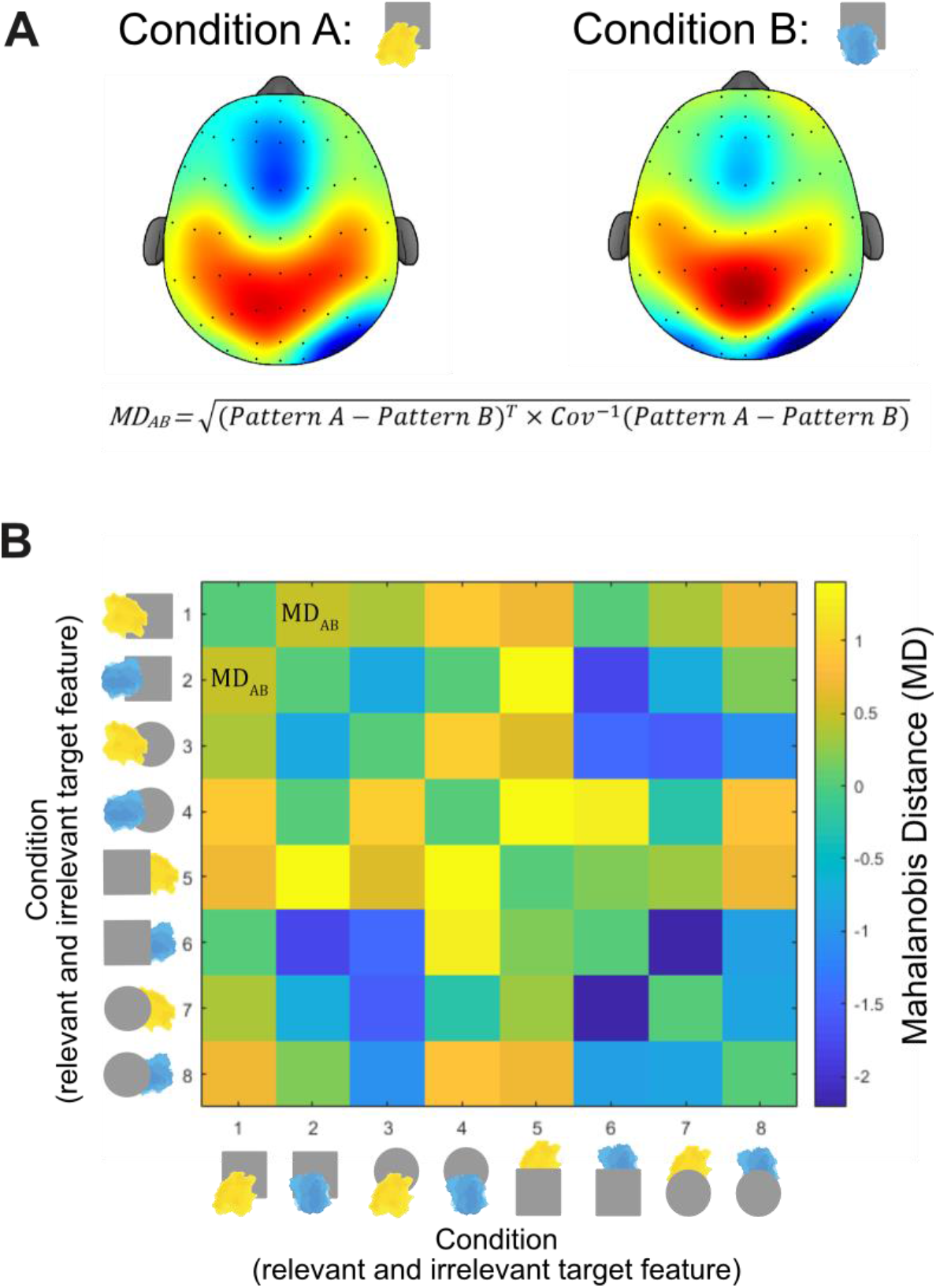
Logic of Representational Similarity Analysis (RSA). A: Trials were divided into conditions based on the relevant and irrelevant features of target stimuli. The dissimilarity between activity patterns was computed as the Mahalanobis distance (MD), which captures the multivariate distance between topographies. B: MDs between conditions are entered into the relevant cells of a representational dissimilarity matrix (RDM). The process outlined in A is repeated until the MDs between each pair of conditions has been computed. The process is then repeated at the next time point, and for subsequent time points of interest. The data RDM produced in B is then regressed against model RDMs that reflect predicted differences in dissimilarity structure for different task variables. These model RDMs are shown in Figure 4.

**Figure 3.**
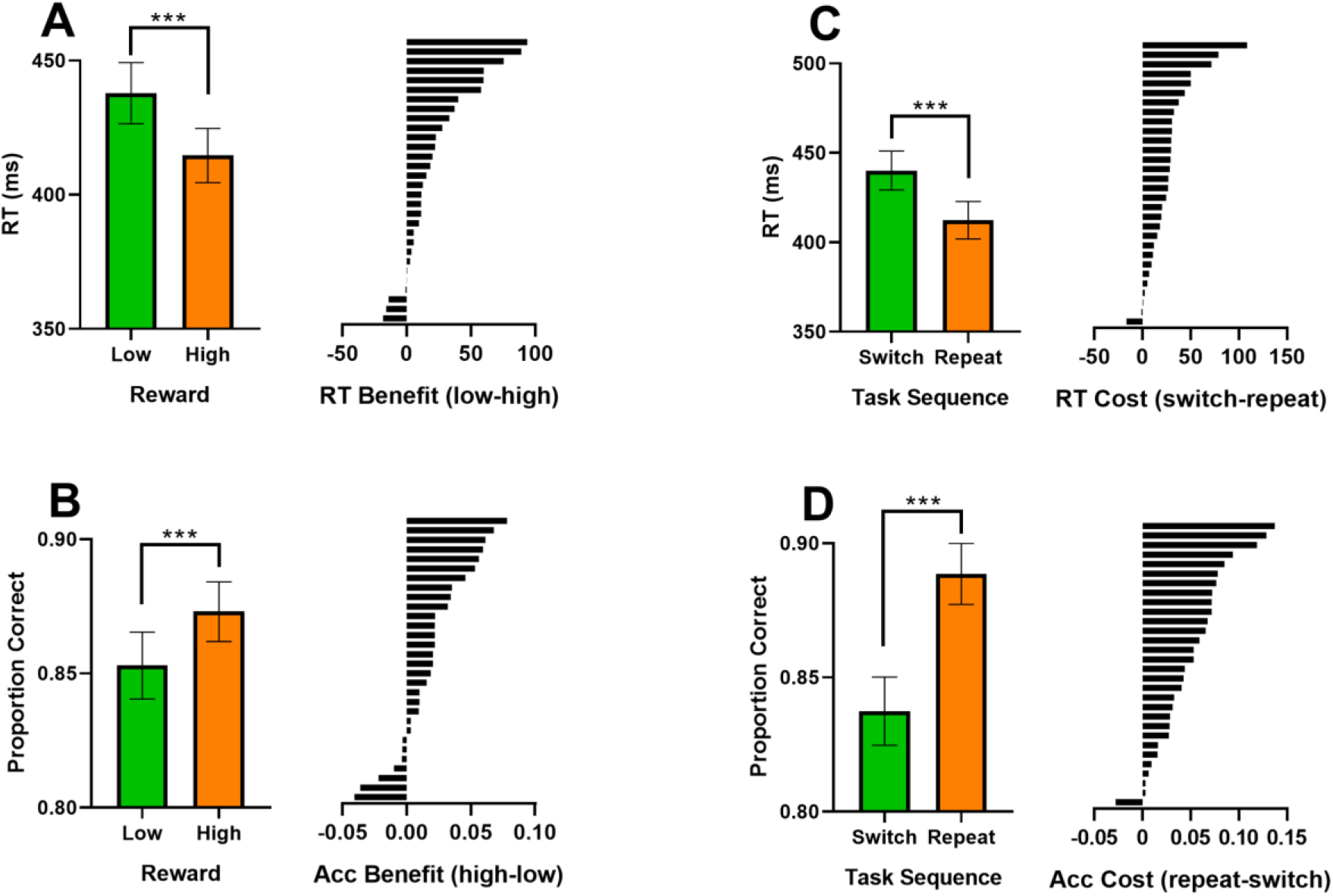
Behavioural performance as a function of reward and task sequence. A: Reaction time for high and low reward trials (left) and the difference between reward conditions for each participant (right). B: Proportion correct (accuracy) for high and low reward trials (left) and the difference between reward conditions per participant (right). C: Reaction time as a function of task sequence (left) and the difference between switch and repeat trials for each participant (right). D: Proportion correct (accuracy) as a function of task sequence (left) and the difference between switch and repeat trials (right) per participant. Error bars show standard error of mean.

##### Neural coding as a function of reward prospect

We repeated the analyses above separately for high and low reward trials. For each participant, trials were divided based on the task-relevant and irrelevant features of the target. Conditions were then averaged over the trial dimension. Mahalanobis distances were calculated between all scalp topographies for the eight conditions, separately at each time point, generating a set of 8×8 RDMs. For these analyses, model RDMs were generated to reflect condition differences based on task coding, the task-relevant feature of the target, the task-irrelevant feature of the target and the motor response. These models followed the same logic as those described in the previous section: zeroes were placed in cells of an 8×8 matrix where two conditions did not differ in the variable of interest (e.g. the task-relevant target feature) and ones were placed in cells where conditions differed in the variable of interest. Model data RDMs were z-scored and entered into a single regression at each time point (including a constant). The regression procedure was performed twice, once using only low reward trials and once using only high reward trials. For this set of analyses, it was particularly important to match the number of high and low reward trials, so that regression coefficients were not higher for high reward conditions because more data was being included in the regression. To address this, these analyses subsampled trials to match the number of high and low reward trials. 100 iterations were performed per participant and final regression coefficients resulted from averaging the estimated regression coefficients over iterations. For task coding, we performed a theory-driven one tailed t-test to assess whether tasking coding was significantly greater under high reward conditions during the pre-target period (averaged from 1400-1800ms). This test was based on Etzel et al. (2015) and remained significant when non-directional. For reward-modulated coding of the relevant target feature, we tested whether the observed results could be driven by correlations with other coding models. To do so, we ran a control analysis in which we first regressed task coding, task-irrelevant feature coding and motor coding against participants’ neural dissimilarity matrices. We then regressed the task-relevant feature coding model against the residual variance, which had not been accounted for by the other models.

##### Neural coding as a function of rule updating

We examined reward-modulated neural coding separately on switch and repeat trials. This was motivated by previous work showing reward prospect benefits switching performance (Shen & Chun, 2011) and can exert a greater performance benefit on switch trials (Kleinsorge & Rinkenauer, 2012), where there is highest interference between task sets and thus where neural separation between task rules could be particularly important in determining behavioural performance. These analyses followed the procedure outlined in the previous section but only used switch or repeat trials, instead of all trials.

##### Neural coding and cognitive performance

To test relationships between neural coding and cognitive performance, we averaged regression coefficients over time windows of interest and correlated them with behavioural scores using non-parametric Spearman correlations. For task coding, we correlated the difference between regression coefficients averaged over the pre-target period (1400-1800) and the difference in switch costs between reward conditions. We undertook the same procedure for reward coding averaged from 0-5000ms from the reward cue. To control for the influence of reward coding when correlating reward-modulated task coding with the changes in performance, we regressed out variance that could be explained by reward coding (averaged from 0-5000ms) from the reward-task coding effect (high-low reward) and the reward-behavioural effect (e.g. RT low reward-RT high reward). We then correlated the residual variance from the reward-task coding effect (averaged over pre-target period: 1400-1800ms or post-target period: 1800-2000ms), with the residual variance in the reward-behavioural effect. We also tested whether the effect of reward on task coding as function of rule updating, was correlated with the difference in switch cost between reward conditions. For this analysis, the interaction term from Figure 6C was averaged over the pre-target period (1400-1800ms).

In addition to time averaged-correlations, we performed time resolved brain-behaviour correlations. To do this, we took participants’ regression coefficients for different task variables at each time point along the trial and correlated this value with their difference in RT (low-high reward) and accuracy (high-low reward). Specifically, these analyses correlated a vector of 30 regression coefficient scores at a single time point with a vector of 30 behavioural scores. The same correlation approach was then performed at the next time point until a correlation had been performed at each time point along the window of interest. The false positive rate for this time-resolved correlation approach was controlled using a cluster-based permutation procedure (detailed in *statistical testing for neural analysis*), in which subjects’ behavioural scores were randomly shuffled over many permutations. With the exception of reward coding (Figure 3C) which used the regression coefficient estimate directly, the neural data for these correlations were differences in regression coefficients between reward conditions. The behavioural data, such as the RT difference between reward conditions, were computed based on all trials used for the behavioural results section. In other words, these behavioural scores did not necessarily use identical trials used in EEG analyses, but instead made use of all behavioural trials available. The correlation analyses performed included neural data reflecting the difference in task coding (Figure 5B), relevant feature coding (Figure 8A), the interaction between reward and relevant feature prioritisation (Figure 8C), as well as motor coding locked to the reward cue (Figure 9A) and the response (Figure 9B). Time-resolved correlations involving reward-differences in neural regression coefficients used the same test windows as their corresponding figure, from which differences were computed. All analyses applied non-parametric Spearman correlations to reduce the influence of outliers in behavioural difference measures, which were more than three scaled median absolute deviations away from the median (RT difference distribution: 2 outliers, lower threshold=−42ms, upper threshold=68ms, median=13ms; accuracy difference distribution: no outliers, lower threshold=−0.06, upper threshold=0.10, median=0.02).

##### Relationships between neural coding of task, sensory and motor information

To test whether reward-induced sensorimotor modulations arose from upstream changes in task coding, we performed a series of correlations between average task and sensorimotor coding regression coefficients. To do so, we selected time windows of interest based on results from the previous analysis section. For task coding, we averaged regression coefficients for each participant from 1400-1800ms. We did not include 1200-1400ms in this average due to a stimulus-evoked signal artefact shown in Figure 4B. For feature coding and motor coding, we averaged regression coefficients from 2000-2400ms, where we observed the peak difference between reward conditions (Figures 5A & 6A). For motor coding locked to the response, we averaged regression coefficients −200-0ms from the response, within which we found the peak difference in motor coding between reward conditions. As initial tests, we correlated average task coding with average feature coding and average motor coding, independent of reward. Significant correlations from this first step were followed up by correlating the mean difference in task coding (high-low reward) with the mean difference in the relevant sensorimotor variable (high-low reward). All correlations were non-parametric Spearman correlations. The Bonferroni-Holm correction was used to correct for multiple comparisons (Holm, 1979). When using this procedure, corrected p-values can become larger than one. To avoid confusion, corrected p-values larger than one were rounded to *p*=1.

**Figure 4.**
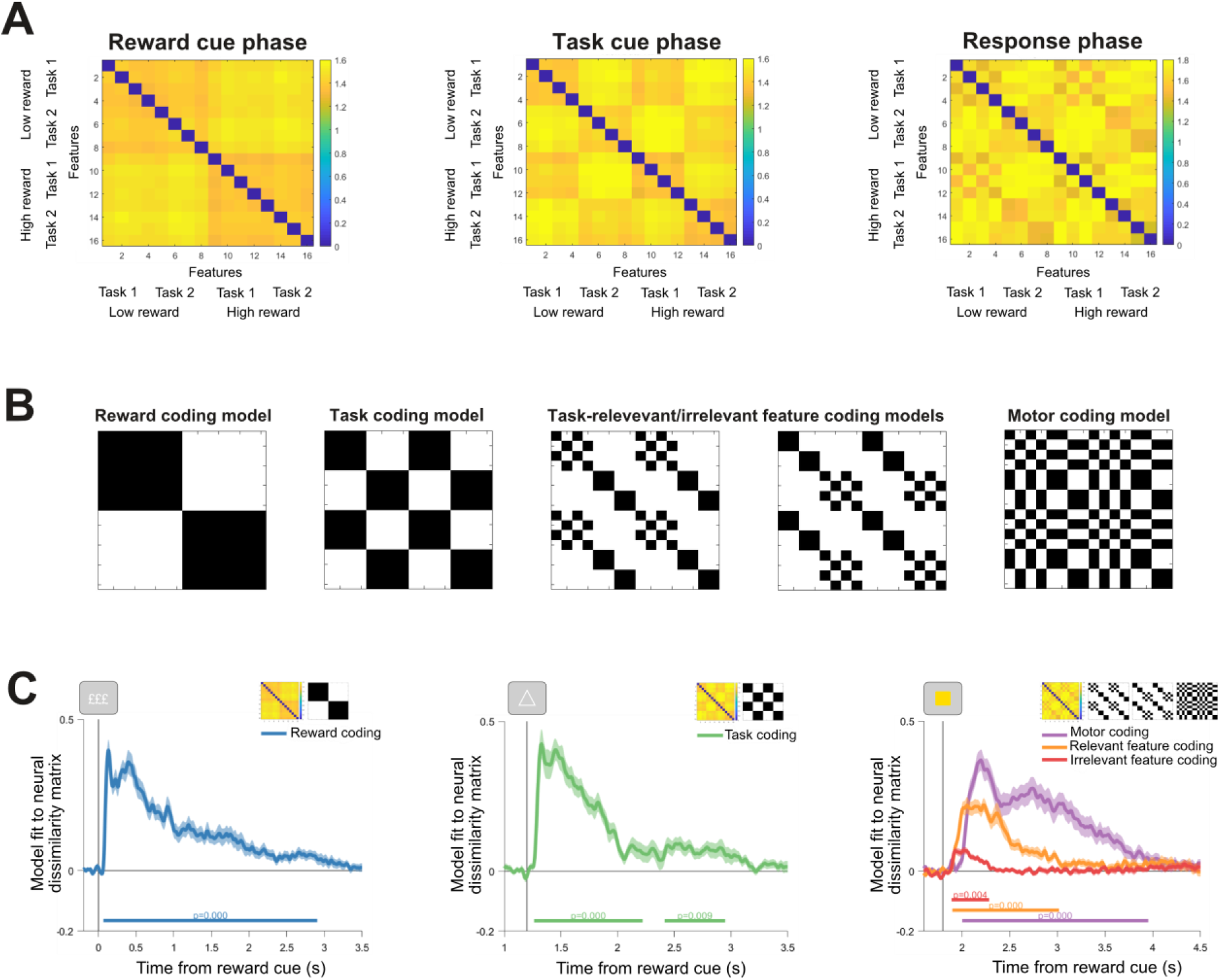
A: Representational dissimilarity matrices averaged across participants and trial periods following: reward cue onset (0-800ms), task cue onset (1200-1800ms) and target onset (1800-2600ms). B: Model representational dissimilarity matrices for: reward coding, task coding, coding of the task-relevant target feature, task-irrelevant target feature and motor coding. C: General linear model regression coefficients from regressing model and neural dissimilarity matrices across time. Shading around principle lines indicates standard error of mean. Cluster-corrected p-values are shown below each time course.

**Figure 5.**
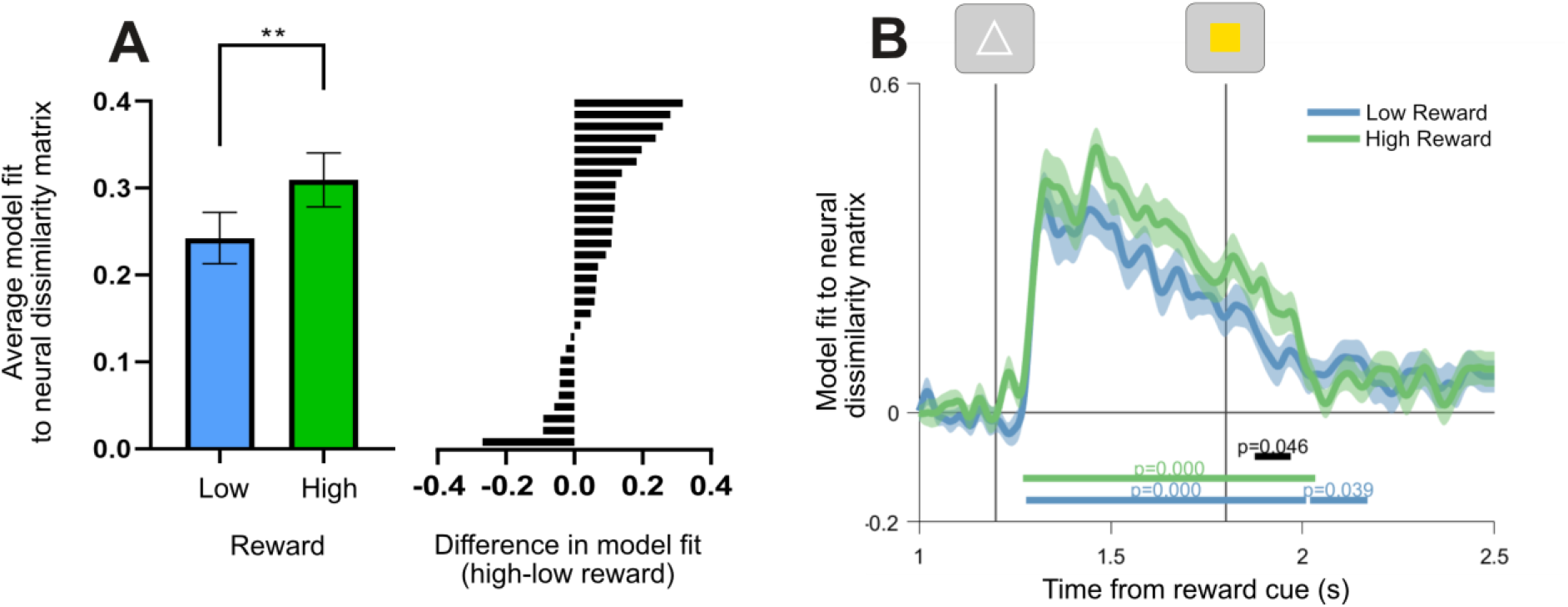
Generalised linear model regression coefficients for task coding as a function of reward. A: coefficients averaged over time points in the pre-target interval (1400-1800ms). The left sub-panel shows group means for average regression betas as a function of reward. The right sub-panel shows the difference between high and low reward regression coefficients for each participant. B: Time resolved regression coefficients for task coding as a function of reward. Vertical lines show onset of the task cue and target respectively. Shading around principle lines indicates standard error of mean. Cluster-corrected p-values are shown below time courses.

**Figure 6.**
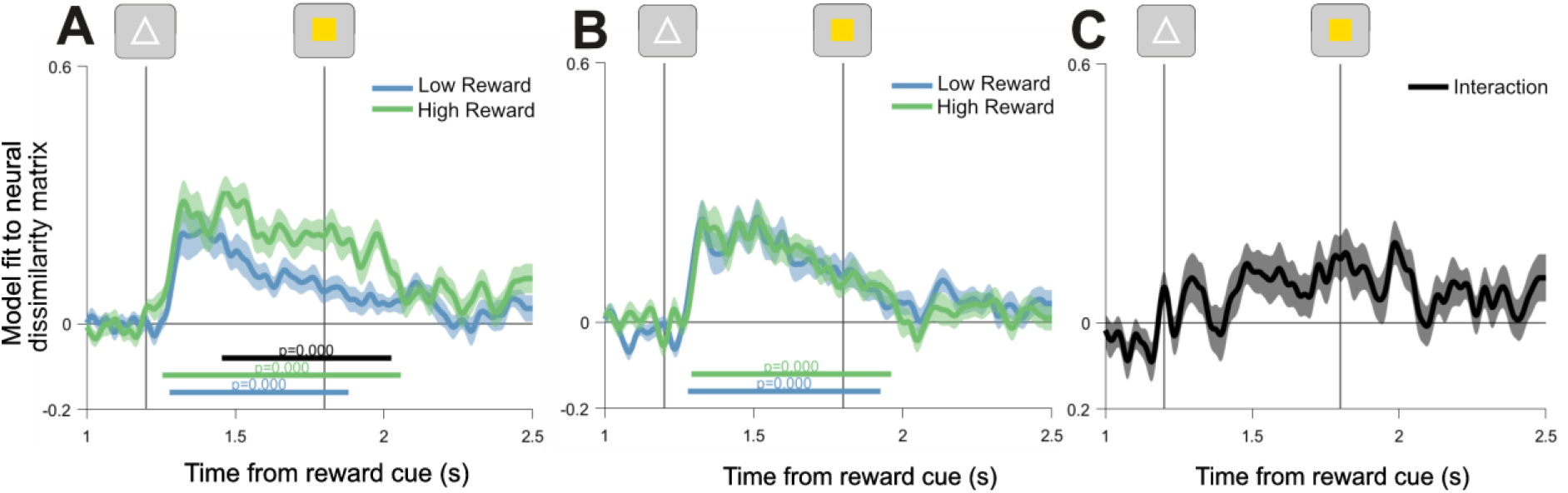
Time resolved regression coefficients for task coding as a function of reward on switch trials (A) and repeat trials (B). C: Interaction between reward and task sequence regression coefficients. This time course is computed by taking the difference between task coding regression coefficients on switch and repeat trials, within each reward condition. Resulting time courses are subtracted to show the extent to which high reward modulates the difference in task coding between switch and repeat trials. Vertical lines show onset of the task cue and target respectively. Shading around principle lines indicates standard error of mean. Cluster-corrected p-values are shown below time courses.

#### Statistical testing for neural analyses

Data were smoothed with a 12ms Gaussian kernel immediately prior to non-parametric cluster-based permutation testing, which was used to correct for multiple comparisons (Maris and Oostenveld, 2007; Sassenhagen & Draschkow, 2018; Spaak, Watanabe, Funahashi, & Stokes, 2017). This non-parametric approach is preferable to parametric statistical alternatives because it does not assume a particular distribution of the data. Cluster-based permutation testing computes a t-statistic at each point in the observed data and many times in permuted data. Each permutation involves shuffling labels for the data, creating a permutated data set that preserves temporal correlations within the EEG signal. Resulting t-statistics are thresholded at an alpha-level of 0.05. Considering only t-values that remain, the cluster of time points with the largest absolute sum of t-statistics (the largest cluster mass) from each permutation is placed into the null distribution. This null distribution shows the likelihood of obtaining a particular cluster mass due to chance. As a final step, candidate clusters from the observed data can then be compared against the null distribution. If the candidate cluster is larger than the 95^th^ percentile of the null distribution, then the effect is considered significant at an alpha level of 0.05. 10,000 permutations were performed to generate null distributions for each analysis in the present study.

## Results

### High reward decreased reaction time and improved accuracy

To assess the impact of reward prospect on behavioural performance, we performed a 2×2 repeated measures ANOVA on RT, with factors of reward (low x high) and task sequence (switch x repeat). This revealed a significant main effect of reward, *F*(1,29)=18.676, *p*<0.001, η2*p*=0.392, which reflected lower RTs on high reward trials (M=415ms, SD=55.45) compared with low reward trials (M=438ms, SD=62.28). There was also a significant effect of task sequence, *F*(1,29)=35.546, *p*<0.001, η2*p*=0.551, driven by lower RTs on repeat trials (M=412ms, SD=57.64) compared with switch trials (M=440ms, SD=59.36). While switch costs were numerically reduced on high compared with low reward trials, the interaction between reward and task sequence did not reach significance, *F*(1,29)=2.662, *p*=0.114, η2*p*=0.084. A control analysis comparing switch trials with repeat trials involving different task cues showed significantly higher RTs on switch trials, confirming the main effect of task sequence was driven by genuine changes in task-set and not simple changes in visual cues (Shapiro-Wilk test for normality: W=0.822, *p*=0.003; Wilcoxon *T*=438.0, p<0.001, matched rank biserial correlation=0.884).

The equivalent 2×2 ANOVA for proportion correct showed the same pattern of results. In particular, the analysis showed a significant main effect of reward, *F*(1,29)=14.268, *p*<0.001, η2*p*=0.330, driven by greater accuracy on high reward trials (M=0.87, SD=0.06) compared with low reward trials (M=0.85, SD=0.07). It also showed a significant main effect of task sequence, *F*(1,29) = 51.894, *p*<0.001, η2*p*=0.642, reflecting greater accuracy for repeat (M=0.89, SD=0.06) compared with switch trials (M=0.84, SD=0.07). We did not detect a significant interaction between reward and task sequence, *F*(1,29)=0.317, *p*=0.578, η2*p*=0.011. Like RT, a control analysis comparing switch trials with repeat trials involving different task cues showed significantly lower accuracy on switch trials, confirming the effect of task sequence on accuracy was driven by genuine changes in task-set (Shapiro-Wilk test for normality: W=0.977, *p*=0.744; t(29)=−6.110, *p*<0.001, *d*=−1.116). To summarise, our behavioural analyses show that high reward prospect improved both RT and accuracy performance.

### Neural coding across the trial

Having established the beneficial impact of reward prospect on cognitive performance, we tested neural coding of task variables across the trial. Reward coding emerged shortly after reward cue onset and was sustained throughout the trial (Figure 4: window tested=0-3500ms from reward cue onset, cluster window=68-2940ms, *p*=0.0002). Task coding peaked shortly after the task rule cue was presented and continued into the response phase (window tested=1200-3500ms from reward cue onset, first cluster=1260-2220ms, first cluster *p*=0.0002, second cluster=2420-2948ms, second cluster *p*=0.0094). Finally, coding of task-relevant and irrelevant target features, as well as motor response coding rose shortly after target presentation (relevant feature coding: window tested = 1800-4500ms, cluster window=1900-3016ms, *p*=0.0002; irrelevant feature coding: window tested= 1800-4500ms from reward cue onset, cluster window=1892-2284ms, cluster *p*=0.0036; motor coding: window tested=1800-4500ms, cluster window=2004-3952ms, cluster *p*=0.0002). In summary, we verified our multivariate analysis approach was sensitive to dynamic temporal changes in neural coding of task variables at sensible stages within the trial.

### Task coding as a function of reward prospect

#### Reward prospect increased proactive task coding

Having verified core task variables were encoded in the EEG signal at the plausible stages within the trial, we tested which variables were influenced by the reward manipulation. We found that average task coding was significantly greater on high reward trials prior to target onset (Figure 5A: window averaged=1400-1800ms; Shapiro-Wilk test for normality: W=0.973, *p*=0.633; theory driven one-tailed test based on Etzel et al. (2015): t(29)=2.881, *p*=0.004, *d*=0.526). Time-resolved permutation analyses confirmed robust encoding of task rules prior to the target under both reward conditions (Figure 5B: window tested=1200-2500ms, first low reward cluster=1280-2008ms, low reward *p*=0.0002; second low reward cluster=2020-2168ms; high reward cluster=1272-2032ms, high reward *p*=0.0002; difference cluster=1876-1968ms, *p*=0.0460).

### Task coding as a function of rule updating

#### Reward-induced increases in task coding were higher on switch trials

Having shown reward prospect modulated task coding overall, we tested whether the effect of reward was amplified during rule updating, where interference between competing rules is highest and thus increased separation between rules could benefit flexible behaviour. This analysis revealed a significant increase in task coding on high reward switch trials compared with low reward switch trials (Figure 6A: window tested=1200-2500, difference cluster=1456-2024ms, *p*=0.0002). By contrast, we did not detect a difference in task coding as a function of reward on repeat trials (Figure 6B: window tested=1200-2500ms, longest candidate cluster=1224-1244ms, *p*=0.6825). The interaction between reward and task coding, as a function of rule updating, was significant when averaged over the pre-target interval (1400-1800ms, *t*(29)=2.9247, *p*=0.0066) and showed a trend towards significance when fully time resolved (Figure 6C: window tested=1200-2500ms, longest candidate cluster=1464-1544ms, *p*=0.0824).

### Task coding and cognitive performance

#### Neural encoding of task rules was associated with performance improvements

To understand the strong performance benefits observed in behaviour, we tested whether task coding was associated with performance improvements (Etzel et al., 2015). The difference in task coding between reward conditions showed a significant correlation with reward-induced changes in RT (Figure 7A: window tested=1200-2500ms, RT difference: cluster window=1860-1968ms, mean Rho=0.4927, *p*=0.0225), but not with accuracy (longest candidate cluster=2280-2320ms, mean Rho=−0.4083, *p*=0.5382). Control analyses confirmed the relationship between reward-induced changes in task coding and RT reflected the difference in RT between reward conditions, rather than either reward condition individually (window tested=1200-1800ms, high reward RT only: longest candidate cluster=1928-1948ms, mean Rho=−0.4004, *p*=0.3030; low reward RT only: longest candidate cluster=1636-1664ms, mean Rho=0.4602, *p*=0.3502). We did not detect significant associations between the reward-rule updating interaction (Figure 6C) and the change in switch costs between reward conditions for RT or accuracy (window averaged=1400-1800, RT switch cost difference: Spearman’s Rho=−0.2725, *p*=0.1449; accuracy switch cost difference: Spearman’s Rho=−0.3351, *p*=0.0703).

**Figure 7.**
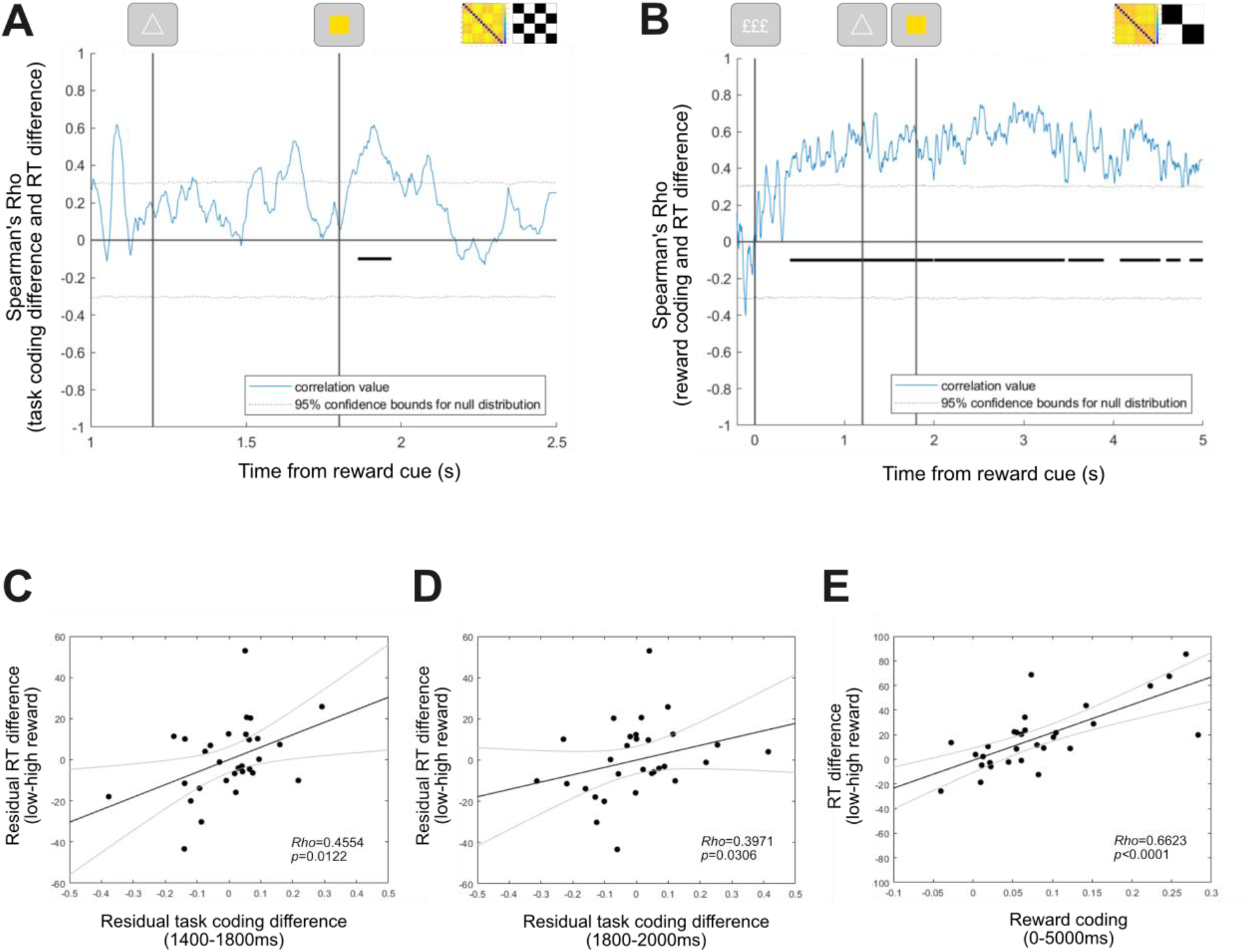
Relationships between reward-modulated task coding and performance. A: Spearman Rho values for correlation between the difference in task coding regression coefficients (high-low reward) and their difference in reaction time between reward conditions. B: Spearman Rho values for correlation between participant reward coding regression coefficients and their difference in reaction time between reward conditions. In both panels A-B, black lines indicate significant correlation clusters, corrected for multiple comparisons using cluster-based permutation testing. Grey dotted lines indicate 95% confidence intervals for the null distribution. C: Scatterplot showing the relationship between the RT difference (low-high reward) and pre-target task coding (high-low reward, 1400-1800ms), after removing variance associated with reward coding. D: Scatterplot showing the relationship between the RT difference (low-high reward) and post-target task coding (high-low reward, 1800-2000ms), after removing variance associated with reward coding. E: Scatterplot showing the relationship between the RT difference (low-high reward) and reward coding (0-5000ms). In panels C-E, black lines indicate linear fits to the data and grey lines indicate 95% confidence intervals of the fits.

Reward coding itself also showed a strong relationship with changes in performance. The magnitude of reward coding was significantly correlated with participants’ difference in RT on high compared with low reward trials (Figure 7A: window tested=0-5000ms, first cluster window=392-1992ms, mean Rho=0.5049, *p*=0.0008; second cluster window=2000-3456ms, mean Rho=0.6001, *p*=0.0004, third cluster window=3496-3892ms, mean Rho=0.5278, *p*=0.0078; fourth cluster window=4072-4524ms, mean Rho=0.5483, *p*=0.0100; fifth cluster=4588-4748ms, mean Rho=0.4487, *p*=0.0415; sixth cluster=4848-4996ms, mean Rho=0.4112, *p*=0.0459). Control analyses correlating reward coding with RT separately on high or low reward trials confirmed this reward coding association was specific to the difference in performance between performance (window tested=0-5000ms, high reward RT only: longest cluster=1188-1248, mean Rho=−0.4113, *p*=0.3258; low reward RT only: longest cluster=4484-4512ms, mean Rho=0.3939, *p*=0.3656). We did not detect a significant correlation between reward coding and the difference in accuracy between reward conditions (window tested=0-5000ms, longest candidate cluster=4-8ms, mean Rho=−0.3822, *p*=0.7373). Similarly, we did not detect significant relationships between reward coding and the difference in switch cost between reward conditions for RT (window averaged=0-5000ms, Spearman’s Rho=0.2912, *p*=0.1185) or accuracy (window averaged=0-5000ms, Spearman’s Rho=0.0472, *p*=0.8045).

To test whether the relationship between reward-driven changes in task coding and improved RT performance was driven by reward coding itself, we ran a control analysis that regressed out variance associated with reward coding (averaged from 0-5000ms) from the reward-task coding effect (high-low) and the reward-RT effect (high-low). We then tested the association between the residual variance in the reward-task coding effect and the residual variance in the reward-RT effect, using non-parametric Spearman correlations. This control indicated that reward-modulated task coding during the pre-target period (averaged from 1400-1800ms) was associated with the change in RT performance, even after removing variance that could be accounted for by reward coding (Figure 7C: Spearman’s Rho=0.4554, *p*=0.0122). This was also the case for reward-modulated task coding during the 1800-2000ms post-target period (Figure 7D: Spearman’s Rho=0.3971, *p*=0.0306). As a final control, we applied this procedure to cluster windows taken from Figures 6A-B (task difference cluster range=1860-1968ms, reward cluster range=392-4996ms). This confirmed the significant time resolved correlation cluster identified for reward-modulated task coding (Figure 6A) was associated with reward-driven changes in RT, independent of reward coding itself (Spearman’s Rho=0.5075, *p*=0.00469).

### Feature coding as a function of reward prospect

#### Reward prospect increased coding of task-relevant target features

After evaluating the impact of reward prospect on task coding, we examined the effect of reward prospect on neural coding of task-relevant and irrelevant target features. Relevant feature coding was observed shortly after target presentation on both high and low reward trials (Figure 8A: window tested=1800-3000ms, low reward cluster=1920-2812ms, *p*=0.0002; first high reward cluster=1916-2892ms, *p*=0.0002). In addition, neural coding for task-relevant features was significantly higher under high reward conditions (window tested=1800-3000ms, first difference cluster=2036-2172, *p*=0.0100; second difference cluster=2252-2404ms, difference *p*=0.0052). A control analysis which regressed the task-relevant feature model against the residual variance, not explained by any of the other coding models, confirmed this result was not driven by correlated regressors (window tested=1800-3000; low reward cluster=1920-2812ms, *p*=0.0002; high reward cluster=1916-2892ms, *p*=0.0002; first difference cluster=2036-2172ms, *p*=0.0120; second difference cluster=2252-2404, *p*=0.0058). Task-irrelevant information was also represented following target onset for both reward levels (Figure 8B: window tested=1800-3000ms, low reward cluster=2008-2284ms, *p*=0.0002; second low reward cluster=2460-2560ms, *p*=0.0470; high reward cluster=1916-2208ms, *p*=0.0008). However, the strength of these coding patterns did not differ as a function of reward (window tested=1800-3000ms, no candidate clusters). As a consequence of these target-related effects, reward prospect showed a significant interaction with the difference in task-relevant and irrelevant feature coding. This reflected a greater difference between task-relevant and irrelevant coding under high reward conditions (Figure 8C: window tested=1800-3000ms, interaction cluster=2240-2364ms, *p*=0.0258). In summary, we found evidence that high reward prospect increased the difference in neural coding for task-relevant and irrelevant target information. This difference was due to a selective increase in the neural coding of task-relevant feature information in high reward contexts.

**Figure 8.**
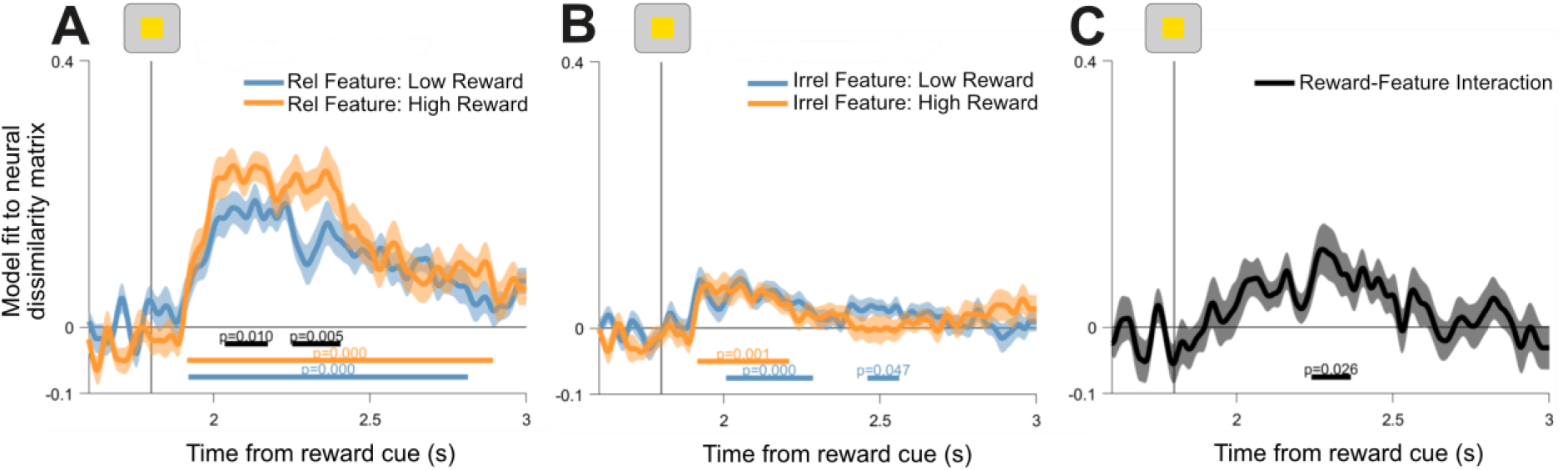
General linear model regression coefficients for coding of target features as a function of reward. A: Task-relevant target feature regression coefficients. B: Task-irrelevant feature regression coefficients. C: Interaction between reward and feature regression coefficients. This time course is computed by taking the difference between task-relevant and task-irrelevant regression coefficients within each reward condition. Resulting time courses are subtracted to show the extent to which high reward increases the difference between task-relevant and irrelevant target features. Vertical lines show target onset. Shading around principle lines indicates standard error of mean. Cluster-corrected p-values are shown below time courses.

### Feature coding as a function of rule updating

#### Reward-feature coding modulations did not differ on switch trials

While high reward prospect increased neural coding of task-relevant feature information overall (Figure 8), the difference in relevant feature coding between reward conditions was not significant when switch and repeat trials were analysed separately (window tested=1800-3000ms, switch trials: longest candidate cluster=2144-2172ms, *p*=0.2274; repeat trials: longest candidate cluster=2104-2148ms, *p*=0.1902).

### Feature coding and cognitive performance

#### Reward-feature coding modulations were not linked to performance improvements

In contrast to the relationships between task coding and performance improvements, we did not detect significant associations between reward-induced changes in feature coding and reward-induced changes in cognitive performance. This was the case for relevant feature coding (window tested=1800ms-3000ms, correlation with RT: longest cluster=1880-1884ms, mean Rho=−0.3809, *p*=0.8172; correlation with accuracy: longest cluster=2660-2676ms, mean Rho=−0.3842, *p*=0.6628), as well as the interaction between reward and relevant feature prioritisation (Figure 8C) (window tested=1800-3000ms, correlation with RT: longest cluster=1868-1880ms, mean Rho=−0.4230, *p*=0.6568; correlation with accuracy: longest cluster=2752-2764ms, mean Rho=0.3789, *p*=0.7073).

### Motor coding as a function of reward prospect

#### Reward prospect increased neural encoding of task-relevant motor output

Having established reward prospect modulated neural activity coding for task-relevant target features, we examined the effect of reward prospect on activity patterns related to the upcoming motor response. When the analysis was locked to the onset of the reward cue, motor coding appeared after target presentation during both high and low reward conditions (Figure 9A: window tested=1800-4500ms, low reward cluster=1996-3716ms, low reward *p*=0.0002; high reward cluster=2016-3792ms, high reward *p*=0.0002) and showed a trend towards higher motor coding under high reward conditions (Figure 9A: window tested=1800-4500ms, difference cluster=2040-2152ms, *p*=0.0900). When the analysis was locked to the onset of the motor response itself, we observed significantly higher motor coding on high reward trials (Figure 9B: window tested =−500-2000ms from response, low reward cluster=−220-1636ms, low reward *p*=0.0002; high reward cluster=−180-1784ms, high reward *p*=0.0002; difference cluster=−36-164ms, *p*=0.0158).

**Figure 9.**
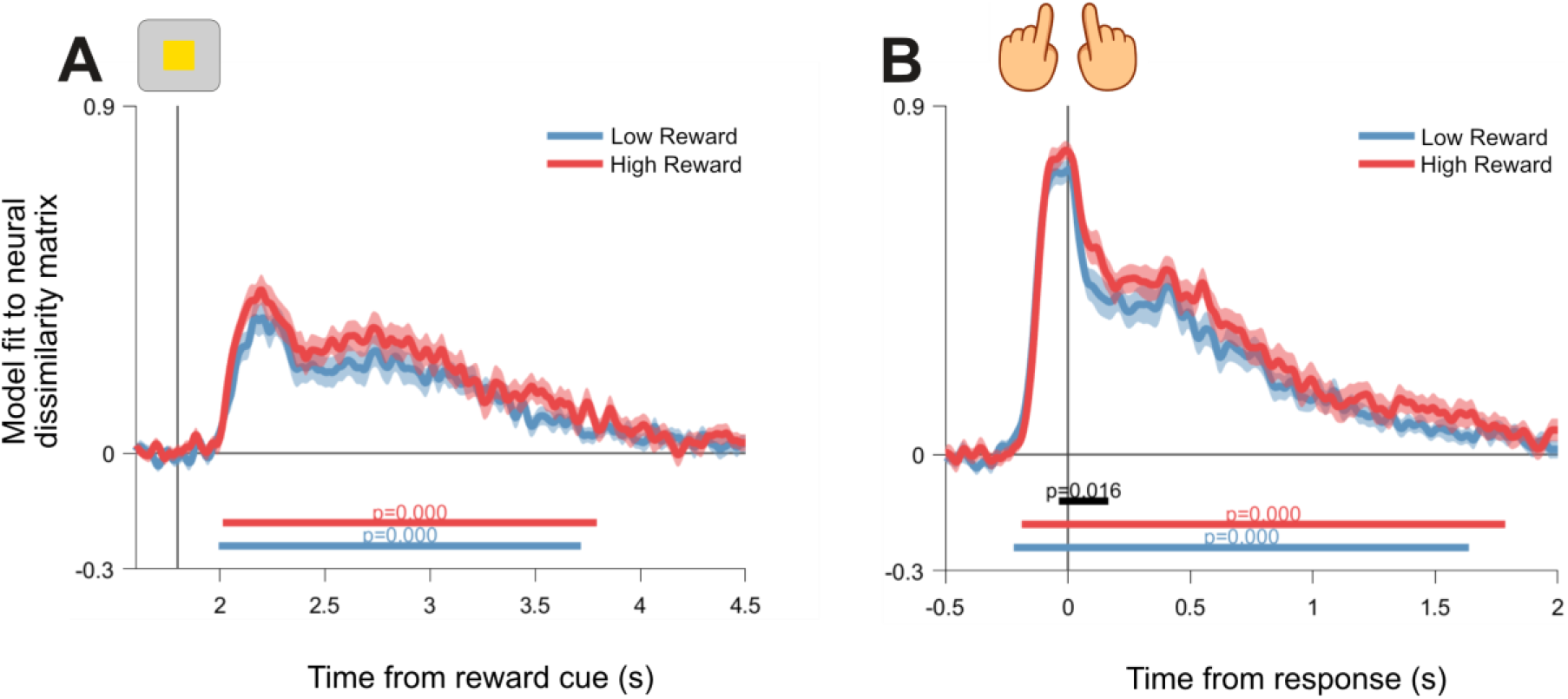
General linear model regression coefficients for coding of the upcoming motor response as a function of reward. A: Motor model regression coefficients for data locked to reward cue onset. The vertical line indicates target onset. B: Motor model regression coefficients for data locked to the response. The vertical line indicates the point responses were made. Shading around principle lines indicates standard error of mean. Cluster-corrected p-values are shown below time courses.

### Motor coding as a function of rule updating

#### Reward prospect modulated motor coding on switch and repeat trials

Based on the reward-motor coding effect observed in our response locked analysis (Figure 9B), we tested whether this effect differed on switch and repeat trials. Motor coding was significantly greater on high compared with low reward switch trials (window tested=−500-2000ms from response, longest difference cluster=−72-180ms, *p*=0.0040), as well as high compared with low reward repeat trials (window tested=−500-2000ms from response, first difference cluster=−84-120ms, *p*=0.0200; second difference cluster=1512-1748ms, *p*=0.0240).

### Motor coding and cognitive performance

#### Reward-motor coding modulations were not linked to performance improvements

When testing whether the reward boost in motor coding (Figure 9B) was associated with changes in performance, we did not detect a significant association between the increase in motor coding (high-low reward) and the change in RT between reward conditions (window tested=−500-2000ms, longest difference cluster=−192 to −152ms, mean Rho=−0.4415, *p*=0.3505). Similarly, we did not detect a significant association between the reward boost in motor coding and the change in accuracy between reward conditions (window tested=−500-2000ms, longest difference cluster=468-524ms, mean Rho=−0.4618, *p*=0.1985).

### Relationships between neural coding of task rule, feature and motor information

#### Reward-induced changes in task coding were not associated with sensorimotor effects

Having established that reward prospect modulated proactive coding of task information, we investigated whether such proactive changes could account for modulations in post-target processing. To do so, we first tested whether average task coding during the pre-target interval (1400-1800ms) was associated with average feature coding (2000-2400ms), as well as average motor coding locked to the reward cue (2000-2400ms) and the response (−200-0ms). Task coding was significantly correlated with coding of the relevant target feature (Spearman’s Rho=0.5355, Bonferroni-Holm corrected *p*=0.0156,) and the difference between task-relevant and irrelevant features (Spearman’s Rho=0.4719, Bonferroni-Holm corrected *p*=0.0455), indicating that participants with greater task coding also tended to exhibit greater prioritisation of task-relevant target features. We did not detect significant correlations between task coding and motor coding locked to the reward cue (Spearman’s Rho=−0.1511, Bonferroni-Holm corrected *p*=1) or the response (Spearman’s Rho=−0.1448, Bonferroni-Holm corrected *p*=1). Having established these relationships, we tested whether reward modulations in task coding during the pre-target period (high-low reward, 1400-1800ms) were correlated with reward modulations in relevant feature coding following target onset (high-low reward, 2000-2400ms). This analysis did not detect a significant correlation between reward-modulated task and reward-modulated feature coding (Spearman’s Rho=0.1097, Bonferroni-Holm corrected *p*=0.8870). The same pattern of results was found when testing the association between reward-modulated task coding (high-low reward, 1400-1800ms) and the extent to which high reward increased the representation of the relevant target feature over the irrelevant feature (2000-2400ms, Spearman’s Rho=0.2957, Bonferroni-Holm corrected *p*=0.4512).

## Discussion

The present study aimed to investigate how rewards modulate neural coding of task rule information to support flexible cognitive control. We employed RSA to examine changes in neural coding of tasks as the prospect of reward changed dynamically, from trial to trial. Using this method, we were able to track neural representations for reward prospect, task rules, task-relevant and irrelevant perceptual features of target stimuli and neural representations related to accurate motor output. We found that high reward prospect boosted the encoding of multiple task variables. Critically, reward increased encoding of the active task rule in preparation for the target and the extent of this increase was associated with reward-based reductions in RT. In addition, reward’s modulatory effect on rule coding was amplified on switch trials, where more flexible processing of rules was needed. Following the target, we observed increased encoding of task-relevant perceptual and motor information under high reward conditions.

Consistent with the results of previous fMRI decoding studies (Etzel et al., 2015; Qiao, Zhang, Chen, & Egner, 2017; Woolgar et al., 2011; Waskom et al., 2014; Wisniewski et al., 2015), RSA successfully tracked neural representations for task rule information. In using this approach, our results replicate the fMRI findings of Etzel et al. (2015) in human electroencephalographic data, showing that high reward prospect increased average task coding prior to target onset, and that the difference in task coding between reward conditions was associated with improvements in cognitive performance. The effect of reward on task coding was most pronounced on switch trials, where task rules needed to be updated relative to the previous trial. This could suggest that dynamic increases in reward prospect primarily promote flexible rule updating, as opposed to maintenance of existing task rule representations in prefrontal regions. Such updating might be mediated by phasic dopamine release in the striatum, which is thought to be important for driving flexible and targeted updating of contextual information in PFC (Westbrook & Braver, 2016; Yee & Braver, 2018). In light of our neural results, showing a larger reward effect on switch trials, it is surprising that there was not a significant corresponding effect in behaviour. We believe this behavioural result should be interpreted with caution, as there was a clear trend in the expected direction, and previous studies have observed robust reward effects on rule switching (Shen & Chun, 2011; Kleinsorge & Rinkenauer, 2012). Overall, the present results provide evidence that high reward prospect can improve cognitive performance by increasing proactive neural representations for task rule information. Moreover, these results seem to be consistent with the theoretical view that reward prospect could help separate the representations of competing task rules, which is especially critical during transitions between different behavioural contexts (Waskom et al., 2014). Indeed such a mechanism could account for the selective effects of reward on switching observed in previous studies (Kleinsorge & Rinkenauer, 2012; Shen & Chun, 2011).

One interesting aspect of these results was that reward-driven changes in task-coding were primarily associated with improvements in RT but not accuracy. This might be viewed as an optimal-control problem, in which subjects maximise expected value over time through balancing speed and accuracy (Bogacz, 2007; Manohar et al., 2015). In our task, accuracy rates were relatively high and while reward significantly increased accuracy, the effect was much smaller than for RT. One possible explanation for this comes from Manohar et al. (2015), who propose an optimal control model to account for the effects of reward on neural noise and behavioural performance. One important prediction of this model is that when signal-to-noise in the neural system is high, rewards will have a disproportionate effect on response vigour and the reduction of RT. These signal-to-noise conditions might well have reflected our task conditions, in which accuracy rates were high and stimuli were unambiguous. A second view comes from the field of reinforcement learning, where it has been proposed that response vigour is calibrated to maximize expected reward rate (Niv, Daw, Joel, & Dayan, 2006) and that this is related to tonic dopamine levels (Beierholm et al., 2013). This view is also compatible with our data, especially given the strong determinant of our RT incentive structure on overall reward rate.

Consistent with the results from previous studies focusing on neural responses to perceptual targets (Padmala & Pessoa, 2011; Serences, 2008; Serences & Saproo, 2010; Hickey & Peelen, 2015), we found that reward enhanced the representation of task-relevant perceptual features. The present results complement this literature in two ways. First, many previous studies have focused on perceptual stimuli associated with reward over many trials (Serences, 2008; Serences & Saproo, 2010; Hickey & Peelen, 2015). Here we show that transient changes in prospective reward can also modulate task-relevant perceptual feature representations. Second, research using prospective reward cues has led to the proposal that reward motivation might benefit attentional filtering, either by enhancing task-relevant perceptual representations or supressing task-irrelevant representations (Pessoa, 2017). Hickey & Peelen (2015) found evidence for both of these mechanisms, showing that reward could either enhance or suppress perceptual representations as a function of task-relevance. In the present study, we report a more selective effect on perceptual representations, wherein high reward prospect increased the neural representation of task-relevant information without impacting the representation of irrelevant information. This might suggest that transient reward coupling with sensory target features has a different impact on perceptual encoding than reward-associations established over many trials.

While task coding and the prioritisation of task-relevant target features were strongly correlated, we did not find evidence that reward-driven modulations in these variables were associated. We are cautious not to over-interpret these null-effects. These results do not rule out the possibility that reward-modulated task coding affects downstream perceptual representations. However, they do raise the possibility of an alternative mechanism, wherein reward prospect acts independently on multiple neural variables. Among these variables, our results suggest that neural encoding of reward prospect and task rule information are important factors associated with dynamic shifts in improving performance. The present results to do not permit conclusive interpretations about the functional role of reward-driven perceptual and motor changes; although we did not detect significant associations between perceptual representations and behavioural measures, this does not imply that these variables were functionally irrelevant to task performance.

How might reward motivation translate into performance improvements more broadly? Previous studies have pointed to the idea that reward motivation might upregulate attention (Etzel et al., 2015; Padmala & Pessoa, 2011; Pessoa & Engelmann, 2010). For instance, reward and attention have been shown to recruit overlapping frontoparietal control regions (Pessoa & Engelmann, 2010) and have analogous effects on electrophysiological signatures of task preparation (van den Berg, Krebs, Lorist, & Woldorff, 2014). One possibility in our study is that reward prospect had additional effects that were not captured by the conditions of our task. For instance, high reward prospect could have increased alertness and temporal attention to information proximal to reward cue presentation. This may explain why associations between behavioural measures and downstream reward effects, such as target feature and motor representations, could then be noisier and less reliable than the strong correlation between behaviour and reward itself.

To conclude, previous work has shown that reward motivation may improve cognitive performance by boosting the neural coding of task rules (Etzel et al., 2015). Here we demonstrate that high reward prospect can increase proactive coding of task rule information and that this proactive effect has a strong relationship with performance improvements. In addition, reward’s effect on rule coding was heightened during switch trials, where task rules were updated relative to the previous trial. This suggests reward prospect might optimise flexible control processes by increasing the neural separation between task rules.

## Acknowledgments

We thank Fabrice Luyckx and Hamed Nili for technical advice, as well as John Grogan and Sanjay Manohar for useful discussion about interpreting the results. This research was funded by a Biotechnology and Biological Sciences Research Council (BB/M010732/1) and James S. McDonnell Foundation Scholar Award (220020405) to Mark G. Stokes, and by the NIHR Oxford Health Biomedical Research Centre. Sam Hall-McMaster is funded by the Rutherford Foundation and William Georgetti Trusts, through the New Zealand Government. Paul S. Muhle-Karbe is funded by the Research Foundation Flanders (grant 12R8817N), the Wellcome Trust (grant 210849/Z/18/Z) and Linacre College Oxford (EPA Cephalosphorin Junior Research Fellowship). Nicholas E. Myers is funded by the Wellcome Trust (201409Z/16/Z) and University College Oxford. The Wellcome Centre for Integrative Neuroimaging is supported by core funding from the Wellcome Trust (203139/Z/16/Z).

